# Gray matter volume in women with the BRCA mutation with and without ovarian removal

**DOI:** 10.1101/2023.11.14.566928

**Authors:** Suzanne T Witt, Alana Brown, Laura Gravelsins, Maria Engström, Elisabet Classon, Nina Lykke, Elisabeth Åvall Lundqvist, Elvar Theordorsson, Jan Ernerudh, Preben Kjölhede, Gillian Einstein

**Affiliations:** Center for Medical Image Science and Visualization (CMIV), Linköping University, Linköping, SWEDEN; BrainsCAN, University of Western Ontario, London, ON, CANADA; Psychology, University of Toronto, Toronto, ON, CANADA; Department of Health, Medicine, and Caring Sciences, Linköping University, Linköping, SWEDEN; Department of Acute Internal Medicine and Geriatrics, and Department of Health, Medicine and Caring Sciences, Division of Prevention, Rehabilitation and Community Medicine, Linköping University, Linköping, SWEDEN; Thematic Studies, Linköping University, SWEDEN; Department of Clinical and Experimental Medicine, Linköping University, Linköping, SWEDEN; Department of Clinical Immunology and Transfusion Medicine, and Department of Biomedical and Clinical Sciences, Linköping University, SWEDEN; Department of Obstetrics and Gynecology and Department of Biomedical and Clinical Sciences, Linköping University, Linköping, Sweden; Tema Genus, Linköping University, Linköping, SWEDEN; Rotman Research Institute, Baycrest Hospital, Toronto, ON, CANADA

## Abstract

**Objective:** Ovarian removal prior to spontaneous/natural menopause (SM) is associated with increased risk of late life dementias including Alzheimer’s disease. This increased risk may be related to the sudden and early loss of endogenous estradiol. Women with breast cancer gene mutations (BRCAm) are counselled to undergo oophorectomy prior to SM to significantly reduce their risk of developing breast, ovarian, and cervical cancers. There is limited evidence of the neurological effects of ovarian removal prior to the age of SM and this is in women without the BRCAm showing cortical thinning in medial temporal lobe structures. A second study in women with BRCAm and BSO noted changes in cognition.

**Methods:** The present study looked at whole-brain changes in gray matter volume using high-resolution, quantitative MRI in women with BRCAm and intact ovaries (BRCA-preBSO) and after surgery with (BSO+ERT) and without (BSO) post-surgical estradiol hormone replacement therapy compared with age-matched women (AMC) with their ovaries.

**Results:** The BRCA-preBSO and BSO groups showed significant gray matter loss in left medial temporal and frontal lobe structures. BSO+ERT exhibited few areas of gray matter reductions compared to AMC. Novel to this study, we also observed that all three BRCAm groups exhibited gray matter increases compared with AMC, suggesting continued plasticity.

**Conclusions:** The present study provides evidence, through gray matter volume reductions, to support both the possibility that the BRCAm, alone, and early life BSO may play a role in increasing the risk for late-life dementia. At least for BRCAm with BSO, post-surgical ERT seems to ameliorate gray matter losses.

## INTRODUCTION

A growing number of studies show that ovarian removal prior to spontaneous (natural) menopause (average age 51) correlates with an increased risk of developing late life dementia and Alzheimer’s disease (AD) ^1–3^. This increased risk is thought to stem from the sudden and early loss of endogenous 17-β-estradiol (E2) ^1,3–6^, where changes in the amount of available E2 have been shown to affect intracellular signaling and neural circuit function in brain regions known to have estrogen receptors (ER) ^7^. A number of post-mortem studies in humans have shown there are ERs in medial temporal lobe structures (including amygdala and hippocampus), entorhinal cortex, and the basal forebrain ^8–12^. Additional post-mortem studies in humans have shown that the dorsolateral prefrontal cortex contains ERs ^12,13^. There is also limited evidence for the presence of ERs in sub-thalamic nuclei but not in other subcortical structures such as putamen, caudate, or globus pallidus ^9,10^. Additionally, there is growing evidence that the cerebellum and its structural development, in particular, are highly sensitive to estrogen ^14^. While the mapping of ER density in the human brain is far from complete, these prior studies give some indication of regions that may be affected both structurally and functionally to the sudden loss of E_2_ caused by ovarian removal. Thus, understanding the earliest brain changes accompanying endogenous E_2_ loss that might lead to increased dementia risk is key to understanding the trajectory towards AD.

Women with BReast CAncer gene mutations, BRCA1 and BRCA2 (BRCAm) are recommended to have prophylactic bilateral salpingo-oophorectomy (BSO; removal of the ovaries and fallopian tubes) prior to spontaneous menopause (SM) to reduce the risk of developing breast and ovarian cancer. Specifically, in women with BRCAm, BSO prior to age 51 has been shown to reduce the risk of developing ovarian, fallopian, and peritoneal cancers by 80% ^15^ and breast cancer by 50% ^16^. Both BRCA1 and BRCA2 are tumor suppressor genes that play important roles in DNA repair ^17–24^. While currently there are no studies investigating how E_2_ loss and the BRCA mutations might intersect to exacerbate neural changes in younger women, we wondered whether carrying BRCAm would, itself, be correlated with changes in regional brain volumes.

To date, there is only limited evidence from two studies examining *in vivo* brain structural changes following BSO in women prior to age 50. Both have findings that suggest brain pathways to dementia. Focusing solely on medial temporal lobe structures previously linked to AD and using a region-of-interest (ROI) approach, women with early life ovarian removal imaged an average of two decades after surgery (average age of 65) had changes in gray matter (GM) thickness and volume previously linked to AD. There were reductions in bilateral amygdalar volume and bilateral parahippocampal thickness but no reduction in bilateral hippocampal volume or cognition ^25^. Interestingly, in this cohort, most of the women were taking conjugated equine estrogens (CEE) with progestins, a much-prescribed form of hormone therapy (HT) in the United States. Since there was no comparator population without HT, it is difficult to know whether HT protected against more changes. The study, however, brought us closer to understanding later life effects of the surgery and perhaps, E2 deprivation. In women with BRCAm and early life BSO, within five years post-BSO (average age 45), the composite hippocampal region, cornu ammonis (CA) 2,3, and dentate gyrus (DG), were reduced in volume ^6^. While these studies point to important late and early regional brain changes, there is still scant information of whole brain changes between six months and two decades post-surgery. The period between these two time points is a critical period to understand since it is when intervention might be most effective. A number of studies have shown that post-surgical estradiol-based therapy (ERT) has a benefit on cognition in both BRCAm carriers (Gervais et al., 2020, 2022) and non-carriers ^26–29^. However, these studies focused solely on behavioral measures of cognition and while one study has shown benefits of ERT on brain structure (Gervais et al., 2022) there is still a lack of evidence as to whether post-surgical ERT has an ameliorative effect on whole brain changes.

Our primary aim therefore was to determine whether women with BRCAm have brain changes within the window between one and ten years post-BSO; our secondary aim was to determine whether simply carrying BRCAm, alone, was associated with brain differences from age matched women without BRCAm and their ovaries, Age Matched Controls (AMC). To do this, we used whole-brain quantitative magnetic resonance imaging (MRI) to examine changes in GM volume in a cohort of healthy Swedish women between the age of 42-54 with BRCAm and BSO prior to the age of 51 within 10 years of surgery comparing them with AMC and BRCA-PreBSO We mapped out GM volume differences between BSO, BSO+ERT, BRCA-PreBSO, and AMC using Synthetic MR ^30^, as this technique has been previously shown to have significantly lower repeat measurement error for GM fraction measured at 3T, making it more sensitive to GM changes than other commonly used methods: SPM, FSL, and Freesurfer (Granberg et al., 2016).

We hypothesized that: 1) BSO would have lower regional GM volumes especially in medial temporal and prefrontal regions compared to BRCA-preBSO and AMC; and 2) BSO+ERT would have regional GM volumes equal to those of BRCA-preBSO and AMC. We also hypothesized that some of the changes would be increased GM volume in regions outside the frontoparietal regions typically associated with executive functions ^32^, suggesting cortical plasticity in service of maintaining appropriate levels of cognitive and executive functioning.

## METHODS

### Study cohort

All participants in this study were drawn from a larger, multi-site longitudinal study on the cognitive and neurological effects of BSO prior to age 50 in Swedish women with BRCAm recruited from local clinics to our study site at Linköping University. All MRI and demographic data included in the current study were drawn from each participant’s first session of this longitudinal study.

Exclusion criteria for all BRCAm women were: 1) being over fifty-five years of age at time of initial consent, 2) contraindications for MRI scanning (e.g., pacemaker, metallic implants, orthodontia, etc.), 3) perimenopause and SM, 4) pregnancy, 5) breastfeeding, 6) chemotherapy, radiation therapy, adjuvant therapy, or hormonal contraception within six months of imaging, 7) untreated health and/or psychiatric conditions, 8) endocrine disorders, 9) and a history of head injury with a loss of consciousness. Since a relatively recent review of chemotherapy-related brain changes in breast cancer survivors noted both GM volume and density losses ^33^ we excluded BRCAm with or without BSO if they reported any history of chemotherapy. Thus, the current study cohort included 20 women with BRCAm with BSO, 10 had BSO and no ERT (average age = 48±4 years, time since BSO = 3±2 years) and 10 had BSO+ERT (average age = 46±3 years, time since BSO = 4±4 years). Nine women with BRCAm without BSO (average age = 39±6 years) were also included as were 10 AMC (average age = 48±3 years) as a referent groups. AMC were recruited from community advertisements. Exclusion criteria for AMC were the same as for BRCAm.

Ethical approval for the study was granted by the Regional Ethical Review Board in Linköping, Sweden (Dnr: 2014/190-31), and all participants provided written consent in accordance with the Helsinki Declaration.

### MRI Scanning

All MRI data were acquired on a Philips 3T Ingenia scanner (Philips Healthcare, Brest, The Netherlands) located at the Center for Medical Image Science and Visualization (CMIV) at Linköping University, Linköping, Sweden. High-resolution 3D T1-weighted images (3D-T1w) were acquired using a standard MPRAGE sequence (FOV 256x240x170; voxel 1x1x1mm^3^; Flip angle 9°; TR/TE = 7/3.2ms; TFE factor 240; Repetitions 1). Quantitative R1-, R2- and proton density (PD)-weighted images were acquired using the QRAPMASTER (quantification of relaxation times and proton density by multi-echo acquisition of a saturation-recovery using turbo spin-echo readout) sequence ^30^, a 2D-FSE multi-dynamic, multi-echo (MDME) saturation recovery spin-echo sequence (TR 4700ms; TE 13/100ms; Flip angle 90°; Refocusing angle 120°; dynamics 4; slices 32; slice thickness (gap) 4(1)mm; voxel size 0.5x0.5mm^2^; SENSE factor 2; acquisition time 6:30). These quantitative images were used to create synthetic T2-weighted (syn-T2w) and GM maps using the SyMRI v8 software (Synthetic MR, Linköping, Sweden; Warntjes et al., 2013).

### Synthetic MR image analysis

Each participant’s syn-T2w image was co-registered to her respective 3D-T1w image. The 3D-T1w image was used to estimate the normalization parameters into Montreal Neurologic Institute (MNI) space, which were then applied to each participant’s GM map. Rather than resampling the GM maps into the standard 1x1x1mm^3^ resolution of the MNI template, the normalized GM map was written out in the original, reconstructed spatial resolution of the quantitative maps (0.5x0.5x5mm^3^). All co-registration and normalization steps were performed in SPM12 (The Wellcome Department of Neuroimaging, UCL, London, UK).

### Statistical analyses

To determine any reductions of GM volume, three independent, between-group t-tests were performed comparing BSO, BSO+ERT, and BRCA-preBSO with AMC. The tests were conducted using the PALM toolbox with 1000 iterations (Winkler et al., 2014), and included age, years of education, and Beck’s Depression Inventory II (BDI-II) scores as covariates of no interest. These three confounds were selected as they have been shown to affect GM volume in previously published population-based studies ^35–37^. Threshold free cluster enhancement (TFCE) was applied to the resulting statistical maps; all imaging results are shown at p < 0.017, FWE (Family Wise Error) corrected for comparing across all voxels and Bonferroni corrected for the three t-tests. To further avoid interpreting spurious clusters, a minimum cluster size of fifty voxels (62.5mm^3^) was applied when reporting significant clusters.

To assess the presence of GM increases, three independent, between-group t-tests were performed comparing BSO, BSO+ERT, and BRCA-preBSO with AMC. Here we also used the PALM toolbox with 1000 iterations was used, and age, years of education, and BDI-II scores were included as covariates of no interest. Threshold free cluster enhancement (TFCE) was applied to the resulting statistical maps, and all imaging results are shown at p < 0.05, FWE (Family Wise Error) corrected for comparing across all voxels. Since this was a secondary aim, no Bonferroni correction was applied to account for conducting the three individual t-tests. However, to avoid interpreting spurious clusters, a minimum cluster size of fifty voxels (62.5mm^3^) was again applied when reporting significant clusters.

Finally, as another exploratory study aim, a whole-brain, one-way four-group ANOVA was performed comparing the GM quantitative maps of the BSO, BSO+ERT, BRCA-preBSO, and AMC. The ANOVA was performed using the PALM toolbox with 1000 iterations ^38^, and included age, years of education, and BDI-II scores as covariates of no interest. Threshold free cluster enhancement (TFCE) was applied to the resulting statistical map, and the whole-brain omnibus F-test result is shown at p < 0.05, FWE (Family Wise Error) corrected for comparing across all voxels. To further avoid interpreting spurious clusters, a minimum cluster size of fifty voxels (62.5mm^3^) was applied when reporting significant clusters in the manuscript and supplemental tables.

## RESULTS

There was a significant difference in age at testing of the participants in the BRCA-preBSO group being, on average, younger than the BSO, BSO+ERT, and AMC. No significant differences were observed for total years of education and Beck’s Depression Inventory II (BDI-II) total score. Additionally, there was no significant difference in the mean time since surgery for the BSO and BSO+ERT.

### Changes in GM volume

#### GM reductions

##### Gray matter reductions in BSO

BSO had significant reductions in GM across the whole brain compared to AMC (Figure 1B, Supplemental Table 2). Within the frontal lobe, reductions were observed in left hemisphere insula, inferior frontal gyrus, precentral gyrus, middle frontal gyrus, inferior orbital frontal gyrus, medial frontal gyrus, superior orbital frontal gyrus, anterior cingulate gyrus, rectal gyrus, and Rolandic operculum. Within the right frontal lobe, reductions were observed in the right anterior cingulate gyrus, middle cingulate gyrus, and medial frontal gyrus. Within the parietal lobe, GM reductions were seen in left inferior parietal lobe and supramarginal gyrus, bilateral precuneus, and right paracentral lobule. Reductions in the temporal lobe were observed primarily in the left hemisphere, including in the superior temporal gyrus, middle temporal gyrus, inferior temporal gyrus, parahippocampus, hippocampus, and amygdala. In the occipital lobe, reductions were noted in bilateral lingual gyrus and right cuneus. Reductions were also observed in the middle and posterior cingulate gyri. Finally, in the cerebellum, reductions were primarily observed in the right hemisphere, including the anterior cerebellum, III, VIII, and the semi-lunar lobule.

**Figure 1:**
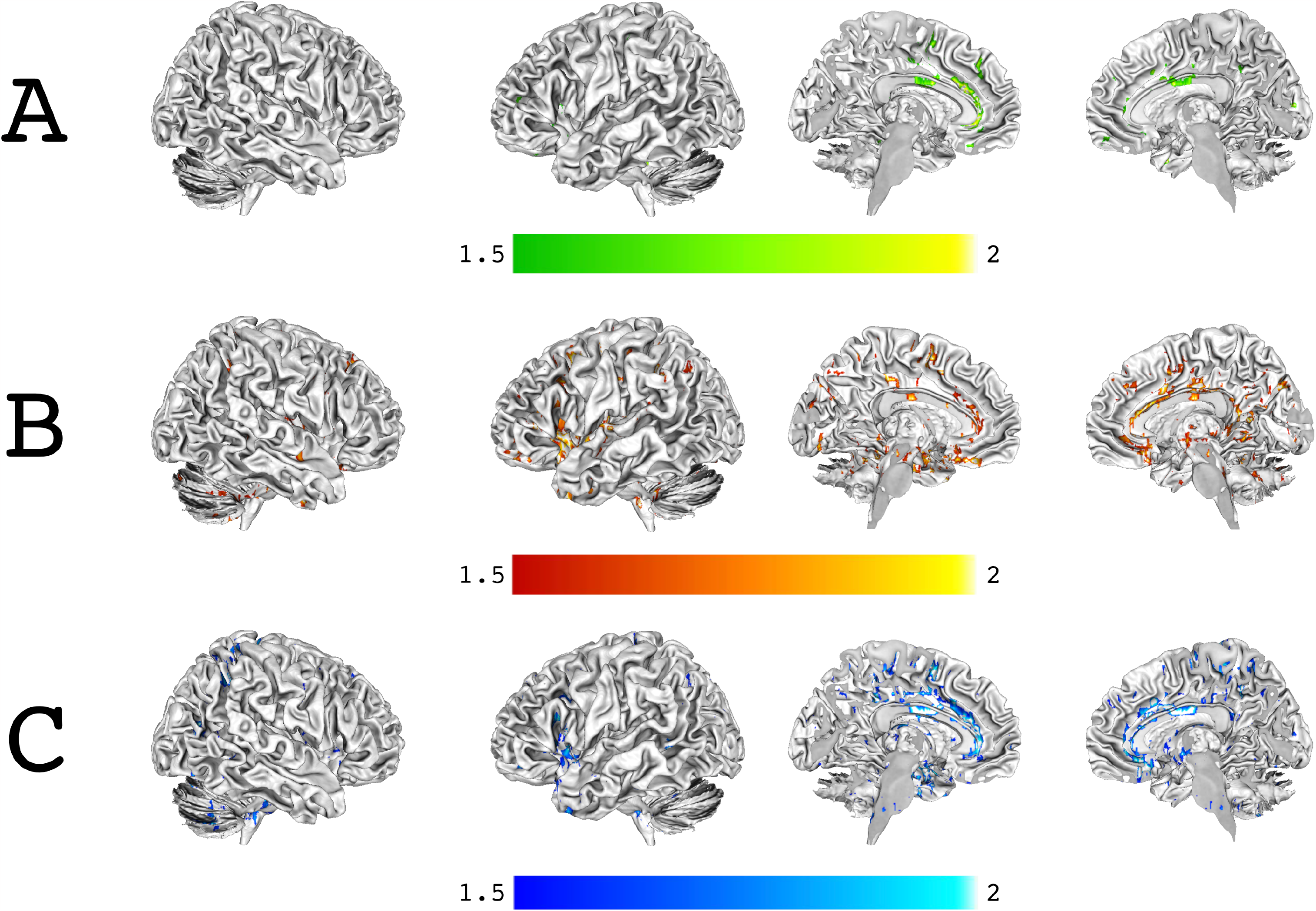
Gray matter reductions compared with age-matched controls. **A:** GM reductions for the BSO+ERT group compared with the AMC group. **B:** GM reductions for the BSO group compared with the AMC group. **C:** GM reductions for the BRCA-preBSO compared with the AMC group. All statistical maps shown at p < 0.0167 FWE corrected across the whole-brain, Bonferonni corrected for the three comparisons. Color bars are scaled in terms of log_10_(p).

##### GM reductions in BSO+ERT

We found few GM reductions in BSO+ERT compared to AMC; these were located in the frontal lobe: left anterior cingulate gyrus, left insula, right middle frontal gyrus, and left rectal gyrus (Figure 1A, Supplemental Table 1).

##### GM reductions in BRCA-preBSO

Surprisingly and contrary to our hypothesis, the BRCA-preBSO group had significant GM reductions across the whole brain similar to the BSO group when compared to AMC (Figure 1C, Supplemental Table 3). In the frontal lobe, reductions were again primarily observed in the left hemisphere, including the anterior cingulate gyrus, precentral gyrus, medial frontal gyrus, middle frontal gyrus, middle orbitofrontal gyrus, insula, and inferior frontal gyrus. Reductions in the right frontal lobe were seen in the anterior cingulate gyrus, medial frontal gyrus, and precentral gyrus. Within in the parietal lobe, GM reductions were observed in the right postcentral gyrus, bilateral precuneus, left paracentral lobule, and right superior parietal lobe.

Within the temporal lobe, the majority of GM reductions were observed in the left hemisphere, including in the parahippocampus, amygdala, hippocampus, uncus, subcallosal gyrus, inferior temporal gyrus, middle temporal gyrus, and superior temporal gyrus. In the right temporal lobe, reductions were noted only in the middle temporal gyrus. Reductions were also observed along the length of the bilateral middle cingulate gyrus. Small areas of GM reductions were also noted in the right lingual gyrus, left caudate, and bilateral cerebellum VIII (Figure 1C, Supplemental Table 3).

#### GM increases

##### Gray matter increases in BSO

Gray matter increases in BSO compared to AMC were observed in a number of select areas across the cortex (Figure 2B, Supplemental Table 5). Within the frontal lobe, increases were noted in bilateral supplementary motor area, bilateral superior frontal gyrus, bilateral precentral gyrus, and the left middle frontal gyrus. In the parietal lobe, increases were seen in the left postcentral gyrus, bilateral superior parietal lobe, and right precuneus. There were also Increases in bilateral middle temporal gyrus and in the occipital lobe in the right middle occipital gyrus and right calcarine fissure.

**Figure 2:**
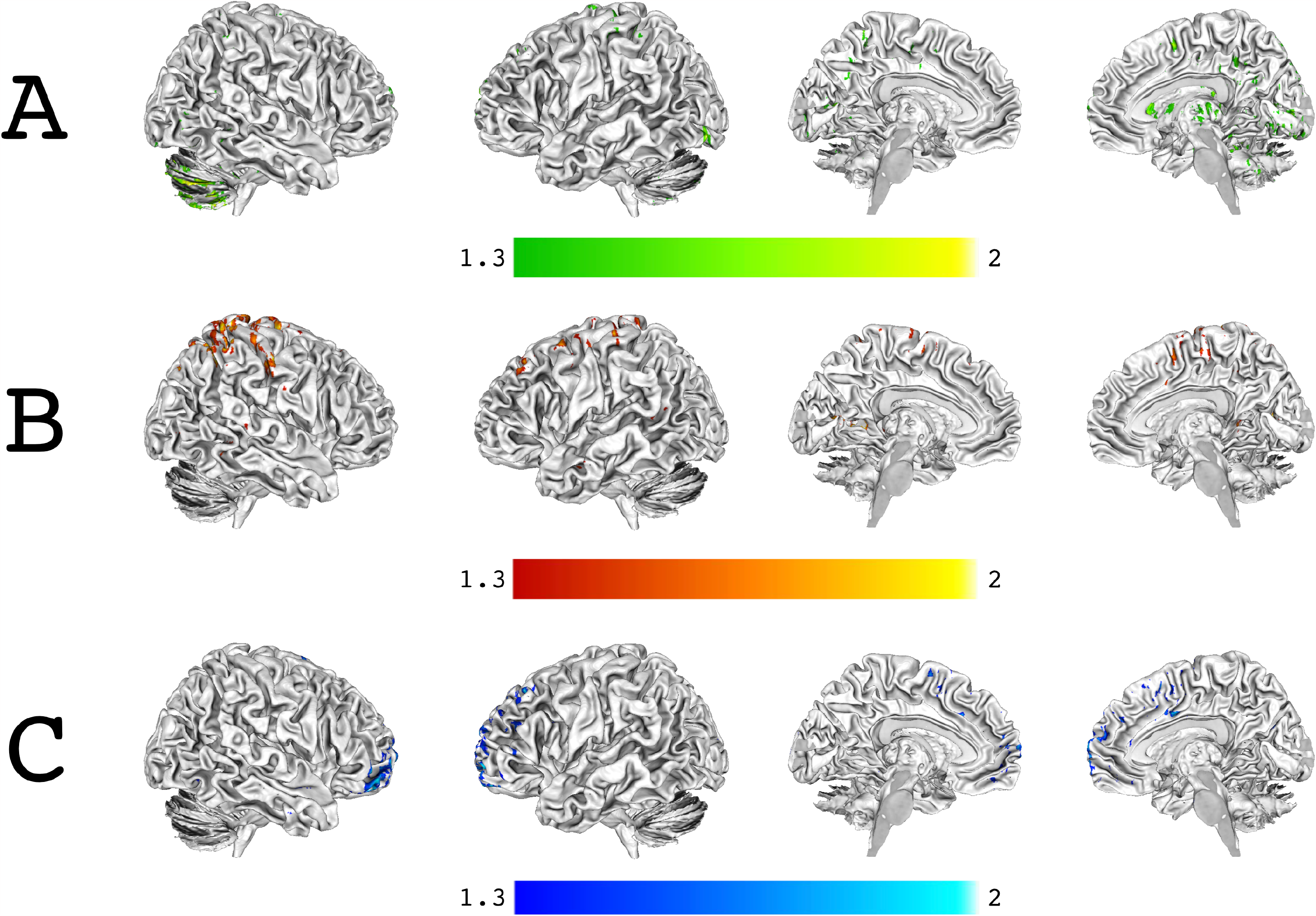
Gray matter increases compared with the age-matched controls. **A:** GM increases for the BSO+ERT group compared with the AMC group. **B:** GM increases for the BSO group compared with the AMC group. **C:** GM increases for the BRCA-preBSO compared with the AMC group. All statistical maps shown at p < 0.05 FWE corrected across the whole-brain. As these statistical analyses were performed in support of our secondary study aim, no Bonferonni correction of the p-values for the three comparisons was performed.

##### GM increases in BSO+ERT

For BSO+ERT, there were numerous areas of GM increases across the cortex, subcortex, and cerebellum compared to AMC (Figure 2A, Supplemental Table 4). In the frontal lobe, increases were observed in bilateral superioral frontal gyrus, bilateral supplementary motor area, and left precentral gyrus. Within the parietal lobe, increases were noted in bilateral precuneus, left postcentral gyrus, left superior parietal lobe, and right inferior parietal lobe. All GM increases in the temporal lobe were observed in the right hemisphere, including the right parahippocampus, right middle temporal gyrus, and right uncus. In the occipital lobe, there were GM increases in the bilateral fusiform gyrus, bilateral calcarine fissure, bilateral inferior occipital lobe, and right lingual gyrus. Subcortically, in right thalamus, right putamen, and right caudate. Finally, cerebellar increases were in bilateral anterior and posterior cerebellum, including Crus1, Crus2, lobe VI, and the declive.

##### Gray matter increases in BRCA-preBSO

The majority of the gray matter increases in the BRCA-preBSO group compared with AMC were observed in the frontal lobe, including bilateral middle frontal gyrus, bilateral superior frontal gyrus, bilateral supplementary motor area, and left inferior frontal gyrus. Smaller areas of increases were noted in the left precuneus, right middle cingulate gyrus, and right cerebellum VIII (Figure 2C, Supplemental Table 6).

### Exploratory aim: Determining differences between BRCA-preBSO and BSO, BSO+ERT, AMC

The results of the exploratory aim are included in the Supplemental Materials (Supplemental Figure 1, Supplemental Table 7). Briefly, GM volume changes were observed across the whole cortex and cerebellum. Notable areas of between group differences were observed across bilateral temporal lobes, including superior, middle, and inferior temporal gyri; bilateral medial temporal structures, including hippocampus, parahippocampus, and amygdala; and bilateral frontal lobe structures, including superior, middle, and inferior frontal gyri and anterior cingulate cortices.

## DISCUSSION

We studied GM volume changes in women with BRCAm both with and without BSO and for the women with BSO both with and without ERT. In line with our hypothesis concerning BSO leading to E2 loss and E2 loss being deleterious to grey matter volume, we observed that compared with the AMC group, the BSO cohort had significant GM reductions in key regions related to late life Alzheimer disease: left temporal lobes, including in hypothesized medial temporal lobe regions including amygdala, hippocampus, and parahippocampus. The BSO+ERT group only had a few, small regions of reduced GM volume, confirming our other hypothesis that post-surgical ERT would have an ameliorative effect but suggesting that ERT does not completely ameliorate brain changes related to BSO.

Overall, our results for the BSO group is in concert with previous studies of older (Zeydan et al., 2018) as well as younger women (Gervais et al., 2022) with BSO prior to age 50; BSO may lead to significant GM reductions in medial temporal lobe structures. As we employed a whole-brain approach, we were further able to refine these findings by demonstrating a strong left-lateralization for the GM reductions in medial temporal structure (amgydala, hippocampus, and parahippocampus). The strong left-lateralization of GM volume reductions observed in the present study is in line with previous findings in mixed sex AD studies, where GM atrophy has been shown to occur earlier and progress more rapidly in the left hemisphere compared with the right hemisphere ^39–42^. This observed asymmetric GM atrophy in the medial temporal lobe has also been observed in normal aging, but a recent study into cortical thinning has shown this asymmetric GM atrophy to be accelerated in AD ^43^. The susceptibility of the left hemisphere in AD has also been demonstrated in mixed sex functional neuroimaging studies, where reductions in activity both during task and rest have been shown to be greater in the left medial temporal structures compared with right medial temporal structures ^44–46^. That these left hemisphere medial temporal GM volume reductions were observed in the BRCA group strongly supports a role for BRCAm, itself, in increasing the risk of developing of late-life dementias. The lack of any medial temporal lobe GM volume reductions in the BSO+ERT group provides additional strong evidence for the role that postsurgical ERT may play in reducing AD risk.

Extending the previous neuroimaging results in humans, we observed that all three study groups exhibited significant GM volume reductions in regions beyond the medial temporal lobe, indicating that the risk of developing late-life dementias conferred by both BRCAm and early life BSO is not contained solely to high ER dense regions, such as the medial temporal lobe. While both BSO and BRCA-preBSO groups had significant GM volume reductions in bilateral middle and inferior frontal regions typically associated with both cognitive and executive functions, the area most consistently found across all three groups was in the anterior cingulate gyrus. Although understudied in dementias, such as AD, mixed sex cohorts show that the anterior cingulate gyrus has degenerative changes in early stage AD, as well as in persons with mild cognitive impairment who go on to convert to AD ^47^. While typically not associated with classic executive functions, such as working memory, the anterior cingulate gyrus is thought to play a role in conflict resolution, salience detection, task selection, and error detection, all cognitive functions necessary in supporting executive functioning ^48^. Further, the anterior cingulate gyrus is known to be structurally connected to insula, prefrontal cortex, amygdala, hypothalamus, and brainstem ^47,48^, suggesting that anterior cingulate gyrus degeneration may result in numerous downstream effects in terms of both brain structure and function.

Novel to this study, we found that all three study groups had significant GM volume increases across the whole brain compared with the AMC group, confirming our hypothesis that all groups would exhibit GM volume increases outside of brain regions typically associated with cognition and executive functions and suggesting continued plasticity. We observed that the BSO, BSO+ERT, and BRCA-preBSO groups also had increased GM volume across the cortex compared with the AMC group, most notably in somatosensory regions and cerebellum. In considering the overall pattern of GM volume reductions across all three study groups, perhaps the most striking finding is that the BSO and BRCA-preBSO groups appear more similar to each other, while the BSO+ERT group more closely resembles the AMC.

Most consistent across all three groups were the increases in sensorimotor regions, including pre- and post-central gyri and the SMA. While the present study cannot provide any specific reason for the significant GM increases in these regions, cortical plasticity seems a likely candidate and the fact that BRCA-preBSO also show it, suggests compensation that might come from detrimental brain effects of the BRCA mutation. As with other brain regions outside the medial temporal lobe, there is only limited evidence in the literature concerning dementia and AD and the sensorimotor cortices, however, at least two mixed sex studies have shown hyperexcitability and increased activation in post-central gyri in mild cognitive impairment and the early stages of AD (Agosta et al., 2010; Ferreri et al., 2016c). The SMA, in particular, is functionally and structurally connected to both medial and lateral prefrontal regions and plays a role in motor planning and learning ^51^. However, the SMA has been more recently shown to be functionally heterogeneous. In addition to its role in planning and execution of movement, several mixed-sex studies have implicated it in working memory ^52^. Focal lesions to SMA are related to working memory deficits, independent of processing speed ^53^. Conversely, greater SMA gyrification is related to working memory improvements ^54,55^, and there is greater SMA engagement with increasing load on the n-back working memory task ^56,57^. The increased GM volume in the SMA observed here may indicate increased engagement of this region in support of working memory functions.

The BSO+ERT group exhibited the largest GM volume increases compared with the AMC group. Of particular note are the GM increases in the right temporal lobe, including medial temporal structures such as the uncus and parahippocampus. A number of mixed sex functional neuroimaging studies have shown increased compensatory activity and hyperexcitability within right-lateralized brain regions, including medial temporal structures, in AD (e.g., Han et al., 2007; Tyrer et al., 2020). That similar right-lateralized GM increases were not observed for either the BSO or the BRCA-preBSO groups, suggests that post-surgical ERT may play a role in this plasticity.

Additionally, volume increases were observed in right subcortical structures, including thalamus, putamen, and caudate, as well as extensively in bilateral cerebellum. Post-mortem ER mapping in humans has indicated that structures like the putamen and caudate do not have high densities of estrogen receptors, suggesting that even without obvious structural changes, the brain may be recruiting those areas less affected by the sudden post-surgical E_2_ loss in an effort to preserve function. However, the same argument cannot be made regarding the increases in the cerebellum, as the cerebellum is known to contain estrogen receptors and be highly sensitive to estrogen ^14^. The cerebellum is known to be highly interconnected with the frontal and prefrontal cortices ^60–62^ and to play a role in supporting a wide array of non-motor functions ^62,63^. A more likely argument for increase GM volume in the cerebellum is that of the brain trying to preserve cognitive function ^61,64^. The GM increases observed here may reflect a compensatory mechanism, as cortical function is lost. However, given that this study solely looked at brain structure and not function, these conclusions are speculative at best.

Perhaps most novel and surprising is that women with the BRCA mutation with intact ovaries and an average of nine years younger, also show decreases in GM volume. This suggests that the BRCA mutation, itself, is detrimental to brain integrity. A number of *in vitro* studies have shown that BRCA1 both 1) accumulates in AD pathologies and tauopathies in general ^17,22,24^ and 2) is generally reduced in patients with AD ^17,20,21,24^. Additionally, reductions of BRCA1 within the brain may be related to learning and memory deficits in mixed-sexed rodents ^21,24^. It is worth noting that 58% (17/29) of our BRCA participants carried BRCA1. While there is less evidence for BRCA2 and dementia, it has been proposed that loss of the BRCA2-mediated DNA repair mechanism may lead to neurodegeneration ^19^. Thus, in addition to increased risk due to the sudden loss of E2, there is evidence from both *in vitro* and *in vivo* rodent and human studies suggesting that carrying BRCAm may further increase the risk of developing late life dementia ^17,20–22,24^. Our results in this regard are very preliminary but support the previous literature on BRCAm and brain changes and raise the interesting possibility that women with the BRCA mutation and BSO may grapple with a double indemnity: issues with gene repair and E2 loss. Interestingly, ERT does help to ameliorate the effects of these two detrimental influences on women’s brain health. Future work is needed to explore the effects of the different types of BRCA mutation as well as the interaction between E2 loss and carrying the mutation.

## LIMITATIONS

This study has a number of limitations, most notably the small cohort size. While the specific SyMRI technique employed is designed to be highly sensitive to GM changes, our conclusions are nonetheless awaiting confirmation in larger studies. One reason underlying the small size was the relatively small number of women recruited to the study who were not prescribed post-surgical ERT. However, even though disaggregating the BSO group into those with and without post-surgical ERT was the primary factor limiting our sample size, we note that this is the first study to specifically examine the structural effects of post-surgical ERT across the whole brain in women with BSO and BRCAm. Further reducing our cohort size was our decision to include only women with no prior reported history of chemotherapy. This decision was made to further limit the number of factors that could confound the GM volume differences we observed. Our small numbers meant that we were also unable to control for BRCAm type. As BRCA1 and BRCA2 are known to influence the brain differently at the cellular level, future studies should examine whether there is an observable difference by BRCA mutation type. Secondly, while we made every effort to match participants across the study groups, there was a significant difference in age, with the BRCA-preBSO group being younger than the other three groups. While it does not seem apparent in the results and age was controlled for in all group-level statistical comparisons, we cannot rule out a small effect of age in our findings. However, it is also worth considering that despite the young age of BRCAm, they had brain changes most similar to those of the women with BSO. Thirdly, while we excluded AMC who reported taking hormonal contraceptives within six months of study enrollment, being a rarer group to recruit, we did not apply this same exclusion criterion to the BRCA-preBSO cohort. Future studies should ensure that women with their ovaries are in known menstrual cycle stages and analyzed by those stages.

## CONCLUSIONS

The present study has demonstrated that even at an early age, shortly after BSO, there is significant evidence, in the form of GM volume reductions, to support the belief that BSO prior to the age of fifty may play a role in increasing the risk for women’s late-life dementia. We also observed evidence to support the role of post-surgical ERT in ameliorating some of these changes. Finally, we provided novel evidence for both the contribution of the BRCA mutation itself as well as likely continued cortical plasticity in all groups but especially in BSO+ERT suggesting that ERT may ameliorate the deleterious effects of BSO.

## ACKNOWLEDGEMENTS

The authors wish to thank Robin Kämpe for his help with data management and transfers.

## FUNDING

Cancerfonden (GE), FORSS (PK), Tema Genus (GE), and the Wilfred and Joyce Posluns Chair in Women’s Brain Health and Aging (GE).

## AUTHOR CONTRIBUTIONS

STW assisted in MRI data collection, performed all image analyses, and drafted the full manuscript. LG and AB helped with formulating the structure of the manuscript and editing the final version. EC helped with data collection. EL, ET, and, JE helped edit the final version of the manuscript. PK obtained support and helped edit the final version of the manuscript. GE conceived of the experiment, obtained support, helped with formulating the structure, and edited the final version.

## SUMMARY SENTENCE

The present study presents evidence of gray matter loss in medial temporal and frontal cortical structures to support the belief that bilateral salpingo-oophorectomy prior to the age of fifty may play a role in increasing the risk for women’s late-life dementia. The study also presents evidence to support the role of post-surgical estrogen replacement therapy in ameliorating some of these changes.

## TABLE CAPTIONS

**Table 1:** Demographic information for the four study groups. The table includes relevant demographic information for the women included in the current study analyses. A significant different in age was observed, with BRCA-preBSO group being younger than the BSO, BSO+ERT, and AMC. No significant differences were observed for total years of education and Beck’s Depression Inventory II (BDI-II) total score. Additionally, there was no significant difference in the mean time since surgery for the BSO and BSO+ERT.

|  | **AMC** | **BRCA** | **BSO** | **BSO-E2** | **P** |
| --- | --- | --- | --- | --- | --- |
| **Number** | 10 | 9 | 10 | 10 |  |
| **Age at scan** | 48.353 | 39.218 | 48.156 | 46.111 | 0.001** |
| **Yrs of education** | 15.75 | 14.667 | 13.25 | 14.35 | 0.14 |
| **BDI-II total** | 4.9 | 8.667 | 8.1 | 7.2 | 0.6 |
| **Time since surgery** | - | - | 3 | 4 | 0.9 |
| **BRCA1** | - | 7 | 4 | 6 |  |
| **BRCA2** | - | 2 | 3 | 3 |  |
| **Cancer** | - | 1 | 2 | 0 |  |
| **Chemotherapy** | - | 0 | 0 | 0 |  |
| **Hormonal contraceptive**<br>**(at time of enrollment)** | 0 | 5 | 0 | 0 |  |

## Notes

CONFLICT OF INTEREST/FINANCIAL DISCLOSURE: None of the authors report any conflicts of interest nor have any financial disclosures related to this manuscript.

### Competing Interest Statement

The authors have declared no competing interest.

