## Supplemental Table 1 for "Gray matter volume in women with the BRCA mutation with and without ovarian removal"

FUNDING: Cancerfonden (GE), FORSS (PK), Tema Genus (GE), and the Wilfred and Joyce Posluns Chair in Women's Brain Health and Aging (GE).

CONFLICT OF INTEREST/FINANCIAL DISCLOSURE: None of the authors report any conflicts of interest nor have any financial disclosures related to this manuscript.

Parts of the data and results of this manuscript have been previously presented at annual meetings for the International Society for Magnetic Resonance in Medicine and the Organization for Human Mappiing.

### TABLES

Table 1: Peak stereotactic coordinates for BSO+ERT gray matter reductions compared with AMC. Coordinates are for all clusters surviving $p < 0.0167$, FWE corrected across the whole brain. All clusters contain at least 50 voxels.

Table 2: Peak stereotactic coordinates for BSO gray matter reductions compared with AMC. Coordinates are for all clusters surviving $p < 0.0167$, FWE corrected across the whole brain. All clusters contain at least 50 voxels.

Table 3: Peak stereotactic coordinates for BRCA-preBSO gray matter reductions compared with AMC. Coordinates are for all clusters surviving $p < 0.0167$, FWE corrected across the whole brain. All clusters contain at least 50 voxels.

Table 4: Peak stereotactic coordinates for BSO+ERT gray matter increases compared with AMC. Coordinates are for all clusters surviving $p < 0.05$, FWE corrected across the whole brain. All clusters contain at least 50 voxels.

Table 5: Peak stereotactic coordinates for BSO gray matter increases compared with AMC. Coordinates are for all clusters surviving $p < 0.05$, FWE corrected across the whole brain. All clusters contain at least 50 voxels.

Table 6: Peak stereotactic coordinates for BRCA-preBSO gray matter increases compared with AMC. Coordinates are for all clusters surviving $p < 0.05$, FWE corrected across the whole brain. All clusters contain at least 50 voxels.

Table 7: Peak stereotactic coordinates for exploratory whole-brain ANOVA results for comparing across all four study groups. Coordinates are for all clusters surviving $p < 0.05$, FWE corrected across the whole brain. All clusters contain at least 50 voxels.

### FIGURES

Figure 1: Exploratory whole-brain ANOVA results for comparing across all four study groups. Statistical map shown at $p < 0.05$ FWE corrected across the whole-brain.

**Table 1:** Peak stereotactic coordinates for BSO+ERT gray matter reductions compared with AMC.

| Region | x | y | z | -log10(p) | voxels | volume (mm³) |
|---|---|---|---|---|---|---|
| | **Frontal Lobe** | | | | | |
| Left Anterior Cingulate Gyrus | -11.5 | 19 | 25 | 2.7 | 528 | 660 |
| Left Insula | -31.5 | 0 | 10 | 2.2 | 73 | 91.25 |
| Right Middle Frontal Gyrus | 45 | 23 | 30 | 2.4 | 65 | 81.25 |
| Left Rectal Gyrus/Medial Frontal Gyrus | -7 | -34 | -25 | 2.4 | 55 | 68.75 |

**Table 2:** Peak stereotactic coordinates for BSO gray matter reductions compared with AMC.

| Region | x | y | z | -log10(p) | voxels | volume (mm$^3$) |
|---|---|---|---|---|---|---|
| | **Frontal Lobe** | | | | | |
| Left Insula/Inferior Frontal Gyrus | -22.5 | 25.5 | -25 | 2.7 | 915 | 1143.75 |
| Left Precentral Gyrus | -42.5 | -7 | 50 | 2.7 | 426 | 532.5 |
| Right Middle Cingulate Gyrus | 2 | -5 | 25 | 2.7 | 404 | 505 |
| Right Anterior Cingulate Gyrus | 9 | 27 | 15 | 2.4 | 379 | 473.75 |
| Left Insula | -31.5 | -17 | 15 | 2.7 | 276 | 345 |
| Left Middle Frontal Gyrus | -37.5 | 6 | 50 | 2.7 | 212 | 265 |
| Left Inferior Orbital Frontal Gyrus/Middle Frontal Gyrus | -40.5 | 38.5 | -10 | 2.7 | 200 | 250 |
| Left Middle Frontal Gyrus | -40 | 15 | 35 | 2.2 | 194 | 242.5 |
| Left Inferior Frontal Gyrus/Middle Frontal Gyrus | -47 | 38.5 | -5 | 2.7 | 192 | 240 |
| Left Medial Frontal Gyrus | -11.5 | 0.5 | 55 | 2.7 | 178 | 222.5 |
| Right Middle Cingulate Gyrus | 8 | 9.5 | 30 | 2.4 | 131 | 163.75 |
| Left Superior Orbital Frontal Gyrus | -21.5 | 55.5 | -5 | 2.4 | 127 | 158.75 |
| Left Rectal Gyrus | -5.5 | 34 | -20 | 2.4 | 119 | 148.75 |
| Left Anterior Cingulate Gyrus | 1 | 34.5 | 5 | 2.7 | 101 | 126.25 |
| Left Precentral Gyrus/BA 6 | -51 | -5.5 | 35 | 2.4 | 97 | 121.25 |
| Right Medial Frontal Gyrus | 2.5 | 21.5 | -20 | 2.4 | 62 | 77.5 |
| Left Rectal Gyrus/Medial Frontal Gyrus/BA 25 | -2.5 | 27.5 | -20 | 2.4 | 54 | 67.5 |
| Left Rolandic Operculum/Precentral Gyrus | -61.5 | -8 | 10 | 2.4 | 53 | 66.25 |
| | **Parietal Lobe** | | | | | |
| Left Inferior Parietal Lobe/Supramarginal Gyrus | -54 | -45 | 35 | 2.7 | 507 | 633.75 |
| Right Precuneus/Posterior Cingulate Gyrus | 7.5 | -46.5 | 15 | 2.4 | 130 | 162.5 |
| Left Precuneus/Superior Occipital Gyrus | -17 | -77.5 | 25 | 2.7 | 126 | 157.5 |

| | | | | | | |
|---|---|---|---|---|---|---|
| Right Parietal Lobe/Paracentral Lobule | 19.5 | -44 | 55 | 2.7 | 85 | 106.25 |
| Right Precuneus | 12 | -52 | 30 | 2.4 | 71 | 88.75 |
| Right Precuneus | 18.5 | -64 | 30 | 2.7 | 57 | 71.25 |
| | **Temporal Lobe** | | | | | |
| Left Superior Temporal Gyrus | -50 | -8.5 | -5 | 2.7 | 561 | 701.25 |
| Left Parahippocampus/Hippocampus/Amygdala | -27.5 | -15 | -25 | 2.7 | 558 | 697.5 |
| Left Superior Temporal Gyrus | -59 | -21.5 | 5 | 2.7 | 329 | 411.25 |
| Left Inferior/Middle Temporal Gyrus | -44 | -0.5 | -40 | 2.7 | 252 | 315 |
| Left Superior Temporal Gyrus | -32 | 14.5 | -40 | 2.4 | 139 | 173.75 |
| Left Middle Temporal Gyrus | -51 | -16.5 | -10 | 2.4 | 104 | 130 |
| Left Parahippocampus/Amygdala/Hippocampus/BA 34 | -16.5 | -3 | -20 | 2.4 | 79 | 98.75 |
| Left Parahippocampal Gyrus | -31 | -34.5 | -10 | 2.4 | 79 | 98.75 |
| Left Superior Temporal Gyrus | -36 | 6.5 | -25 | 2.4 | 76 | 95 |
| Right Inferior Temporal Gyrus | 45.5 | -1.5 | -40 | 2.4 | 73 | 91.25 |
| Left Temporal Lobe/Parahippocampal Gyrus | -23 | 2.5 | -30 | 2.4 | 71 | 88.75 |
| | **Occipital Lobe** | | | | | |
| Right Cuneus/Precuneus | 21.5 | -72.5 | 20 | 2.7 | 787 | 983.75 |
| Left Lingual Gyrus/Parahippocampal Gyrus | -8.5 | -38 | 5 | 2.7 | 208 | 260 |
| Right Cuneus | 12 | -87 | 20 | 2.7 | 137 | 171.25 |
| Right Lingual Gyrus/Parahippocampal Gyrus | 22 | -51 | 0 | 2.2 | 54 | 67.5 |
| | **Cingulate Gyrus** | | | | | |
| Left Middle Cingulate Gyrus/BA 31 | -11.5 | -32 | 40 | 2.7 | 463 | 578.75 |
| Right Posterior Cingulate Gyrus | 13 | -56.5 | 10 | 2.7 | 80 | 100 |
| Left Cingulate Gyrus/Paracingulate Gyrus | -6.5 | 9.5 | 45 | 2.4 | 52 | 65 |

| | Cerebellum | | | | | |
|---|---|---|---|---|---|---|
| Left Anterior Cerebellum/Culmen | 3.5 | -47 | -10 | 2.7 | 142 | 177.5 |
| Right Anterior Cerebellum | 4.5 | -47.5 | -20 | 2.4 | 72 | 90 |
| Right Cerebellum III | 19 | -28.5 | -35 | 2.1 | 70 | 87.5 |
| Right Cerebellum semi-lunar lobule | 9 | -68 | -50 | 2.4 | 66 | 82.5 |
| Right Cerebellum VIII | 25 | -55.5 | -60 | 2.2 | 60 | 75 |

**Table 3:** Peak stereotactic coordinates for BRCA-preBSO gray matter reductions compared with AMC.

| Region | x | y | z | -log10(p) | voxels | volume (mm³) |
|---|---|---|---|---|---|---|
| | **Frontal Lobe** | | | | | |
| Right Anterior Cingulate/BA 24/ Medial Frontal Gyrus | 6.5 | 22.5 | -20 | 2.7 | 2843 | 3553.75 |
| Left Anterior Cingulate Gyrus | -7.5 | 35.5 | 10 | 2.7 | 415 | 518.75 |
| Left Precentral Gyrus | -29 | -25 | 45 | 2.4 | 259 | 323.75 |
| Left Medial Frontal Gyrus/Cingulate Gyrus | -12 | 1 | 50 | 2.7 | 247 | 308.75 |
| Left Middle Frontal Gyrus | -38 | 10.5 | 35 | 2.4 | 225 | 281.25 |
| Right Medial Frontal Gyrus/BA 6 | 2.5 | -25 | 60 | 2.7 | 138 | 172.5 |
| Left Inferior Operculum/Precentral Gyrus | -50 | 11 | 10 | 2.7 | 117 | 146.25 |
| Left Middle Orbitalfrontal Gyrus | -33.5 | 51 | -5 | 2.7 | 88 | 110 |
| Left Insula | -30.5 | 26.5 | 5 | 2.4 | 88 | 110 |
| Left Inferior Frontal Gyrus | -50.5 | 23.5 | 15 | 2.4 | 84 | 105 |
| Left Inferior Frontal Gyrus | -46.5 | 17 | 0 | 2.7 | 62 | 77.5 |
| Right Medial Frontal Gyrus | 6.5 | -18.5 | 60 | 2.2 | 62 | 77.5 |
| Left Precentral Gyrus | -27.5 | -27.5 | 65 | 2.4 | 61 | 76.25 |
| Left Middle Frontal Gyrus/ Inferior Orbitalfrontal Gyrus | -40.5 | 42 | -5 | 2.2 | 58 | 72.5 |
| Right Precentral Gyrus/Postcentral Gyrus | 46.5 | -5.5 | 30 | 2.1 | 56 | 70 |
| Left Inferior Frontal Gyrus | -39 | 25 | 0 | 2.7 | 52 | 65 |
| | **Parietal Lobe** | | | | | |
| Right Parietal Lobe | 35 | -46 | 35 | 2.4 | 182 | 227.5 |
| Left Precuneus/Paracentral Lobule | -5 | -43.5 | 50 | 2.4 | 145 | 181.25 |
| Right Parietal Lobe/Postcentral Gyrus | 8 | -33 | 70 | 2.4 | 122 | 152.5 |
| Right Precuneus/Parietal Lobe | 8.5 | -48 | 45 | 2.7 | 120 | 150 |
| Left Paracentral Lobule/Precuneus | -1 | -44 | 60 | 2.4 | 51 | 63.75 |

| | | | | | | |
|---|---|---|---|---|---|---|
| Right Superior Parietal Lobe/BA 7 | 14 | -57.5 | 60 | 2.2 | 50 | 62.5 |
| | **Temporal Lobe** | | | | | |
| Left Parahippocampus/Amygdala/ Hippocampus/Uncus | -19 | -14 | -20 | 2.7 | 1236 | 1545 |
| Left Parahippcampal Gyrus | -15.5 | -23 | -15 | 2.7 | 116 | 145 |
| Left Subcallosal Gyrus | -7.5 | 4 | -20 | 2.2 | 81 | 101.25 |
| Left Inferior Temporal Gyrus/Fusiform Gyrus | -43.5 | -8 | -30 | 2.4 | 79 | 98.75 |
| Left Inferior Temporal Gyrus/BA 20 | -53.5 | -28 | -20 | 2.2 | 77 | 96.25 |
| Left Superior Temporal Gyrus | -45 | -52.5 | 15 | 2.7 | 75 | 93.75 |
| Right Middle Temporal Gyrus | 53 | -64.5 | 10 | 2.2 | 66 | 82.5 |
| Left Middle Temporal Gyrus | -62 | -47.5 | 0 | 2.7 | 60 | 75 |
| Left Superior Temporal Gyrus | -54.5 | 5.5 | -5 | 2.4 | 56 | 70 |
| Left Parahippcampal Gyrus | -28 | -11.5 | -25 | 2.1 | 55 | 68.75 |
| | **Cingulate Gyrus** | | | | | |
| Left Cingulate Gyrus/Middle Cingulate Gyrus | -11 | -17 | 35 | 2.7 | 826 | 1032.5 |
| Right Cingulate Gyrus/Middle Cingulate Gyrus/ Anterior Cingulate Gyrus | 10 | 19 | 35 | 2.7 | 321 | 401.25 |
| Right Middle Cingulate Gyrus | 12 | -34 | 40 | 2.7 | 204 | 255 |
| Right Cingulate Gyrus | 14 | -10.5 | 30 | 2.7 | 162 | 202.5 |
| Left Cingulate Gyrus/Middle Cingulate Gyrus | -7.5 | -38.5 | 35 | 2.4 | 161 | 201.25 |
| Left Cingulate Gyrus/Middle Cingulate Gyrus/ Anterior Cingulate Gyrus | -10.5 | 19.5 | 25 | 2.2 | 132 | 165 |
| Right Cingulate Gyrus/Middle Cingulate Gyrus | 12.5 | 1.5 | 40 | 2.1 | 67 | 83.75 |
| | **Occipital Lobe** | | | | | |
| Right Lingual Gyrus | 17.5 | -84 | -10 | 2.2 | 81 | 101.25 |
| | **Subcortex** | | | | | |

| | | | | | | |
|---|---|---|---|---|---|---|
| Left Caudate | -11 | 5.5 | 15 | 2.7 | 56 | 70 |
| | **Cerebellum** | | | | | |
| Right Cerebellum VIII | 16 | -66.5 | -55 | 2.7 | 236 | 295 |
| Left Cerebellum VIII | -28.5 | -48 | -50 | 2.2 | 81 | 101.25 |

**Table 4:** Peak stereotactic coordinates for BSO+ERT gray matter increases compared with AMC.

| Region | x | y | z | -log10(p) | voxels | volume (mm³) |
|---|---|---|---|---|---|---|
| | **Frontal Lobe** | | | | | |
| Right Superior Frontal Gyrus/BA 10 | 15 | 70.5 | 20 | 2.1 | 714 | 892.5 |
| Right Superior Frontal Gyrus | 22.5 | 56 | 40 | 2.1 | 202 | 252.5 |
| Right Superior Frontal Gyrus | 22 | 50 | 45 | 2.2 | 198 | 247.5 |
| Left Superior Frontal Gyrus | -23.5 | 66 | 20 | 2 | 117 | 146.25 |
| Left Superior Frontal Gyrus | -11.5 | 69 | 15 | 1.9 | 98 | 122.5 |
| Right Supplementary Motor Area/ Medial Frontal Gyrus | 5.5 | 3.5 | 55 | 1.9 | 60 | 75 |
| Left Precentral Gyrus | -19.5 | -11.5 | 75 | 1.6 | 60 | 75 |
| Left Superior Frontal Gyrus | -20 | 68 | 5 | 1.5 | 52 | 65 |
| Left Supplementary Motor Area | -3.5 | 5.5 | 50 | 1.4 | 50 | 62.5 |
| | **Parietal Lobe** | | | | | |
| Left Precuneus | -5.5 | -48.5 | 65 | 1.8 | 301 | 376.25 |
| Right Precuneus | 7 | -44.5 | 40 | 1.7 | 170 | 212.5 |
| Left Precuneus | -1.5 | -58 | 35 | 1.7 | 153 | 191.25 |
| Left Superior Parietal Lobe | -29.5 | -63.5 | 45 | 1.8 | 80 | 100 |
| Left Postcentral Gyrus/Precentral Gyrus | -36 | -29.5 | 70 | 1.7 | 78 | 97.5 |
| Right Precuneus | 4.5 | -57.5 | 30 | 1.5 | 74 | 92.5 |
| Right Inferior Parietal Lobe | 55.5 | -42.5 | 50 | 1.7 | 64 | 80 |
| Left Postcentral Gyrus | -26.5 | -34 | 45 | 1.6 | 56 | 70 |
| Right Precuneus | 9.5 | -80 | 55 | 1.6 | 56 | 70 |
| Left Postcentral Gyrus | -31 | -37.5 | 55 | 2 | 53 | 66.25 |
| Right Inferior Parietal Lobe/BA 40 | 48.5 | -43 | 60 | 1.6 | 50 | 62.5 |
| | **Temporal Lobe** | | | | | |

| | | | | | | |
|---|---|---|---|---|---|---|
| Right Parahippocampal Gyrus | 24 | -45.5 | -10 | 1.8 | 280 | 350 |
| Right Middle Temporal Gyrus | 48.5 | -70 | 0 | 1.7 | 157 | 196.25 |
| Right Parahippocampal Gyrus | 23.5 | -21 | -30 | 1.7 | 131 | 163.75 |
| Right Uncus/BA 20 | 29.5 | -15 | -35 | 1.5 | 78 | 97.5 |
| Right Middle Temporal Gyrus | 58.5 | -40.5 | -5 | 1.5 | 71 | 88.75 |
| | **Occipital Lobe** | | | | | |
| Left Fusiform Gyrus/Lingual Gyrus | -25.5 | -82 | -5 | 2 | 535 | 668.75 |
| Right Calcarine Fissure | 10 | -84.5 | 5 | 2.1 | 443 | 553.75 |
| Left Inferior Occipital Lobe | -41.5 | -83 | -10 | 2.1 | 410 | 512.5 |
| Right Fusiform Gyrus | 35.5 | -58.5 | -20 | 2.1 | 273 | 341.25 |
| Left Calcarine Fissure | -18 | -74.5 | 10 | 2.1 | 217 | 271.25 |
| Right Calcarine Fissure/Cuneus | 12.5 | -71.5 | 10 | 2.4 | 171 | 213.75 |
| Right Lingual Gyrus | 20 | -64.5 | -10 | 1.9 | 170 | 212.5 |
| Right Inferior Occipital Gyrus | 31.5 | -88.5 | -10 | 2.1 | 73 | 91.25 |
| Right Fusiform Gyrus | 47 | -59 | -20 | 1.5 | 69 | 86.25 |
| | **Subcortex** | | | | | |
| Right Thalamus/Hippocampus/ Parahippocampal Gyrus | 20.5 | -14.5 | -5 | 2.4 | 1891 | 2363.75 |
| Right Putamen | 24 | 4 | 0 | 2.1 | 293 | 366.25 |
| Right Thalamus | 7.5 | -17 | 5 | 1.7 | 149 | 186.25 |
| Right Caudate | 9 | 9.5 | 5 | 1.6 | 93 | 116.25 |
| | **Cerebellum** | | | | | |
| Right Posterior Cerebellum/ Crus1/Crus2 | 39 | -53.5 | -60 | 2.7 | 5270 | 6587.5 |
| Right Cerebellum Crus2 | 2.5 | -83.5 | -25 | 2.1 | 465 | 581.25 |
| Right Posterior Cerebellum | 5 | -62 | -40 | 1.7 | 259 | 323.75 |

| | | | | | | |
|---|---|---|---|---|---|---|
| Left Posterior Cerebellum/VI | -4.5 | -74 | -20 | 1.7 | 245 | 306.25 |
| Right Anterior Cerebellum/Culmen | 11.5 | -41.5 | -25 | 1.8 | 220 | 275 |
| Left Cerebellum Crus2 | -5 | -81 | -35 | 1.9 | 208 | 260 |
| Right Cerebellum VI | 20.5 | -71 | -15 | 2.1 | 163 | 203.75 |
| Right Vermis 4-5 | 4 | -54.5 | 0 | 2.1 | 160 | 200 |
| Left Posterior Cerebellum | -2 | -66.5 | -25 | 1.7 | 149 | 186.25 |
| Left Anterior Cerebellum | -3.5 | -64 | -10 | 1.7 | 142 | 177.5 |
| Right Cerebellum/Culmen | 29.5 | -43.5 | -25 | 1.9 | 110 | 137.5 |
| Left Anterior Cerebellum | -4 | -56.5 | -35 | 1.8 | 85 | 106.25 |
| Right Cerebellum VI | 22 | -57 | -25 | 1.6 | 79 | 98.75 |
| Right Posterior Cerebellum/Declive | 33.5 | -67 | -30 | 1.5 | 60 | 75 |
| Left Posterior Cerebellum/Declive | -45 | -53.5 | -25 | 1.9 | 59 | 73.75 |
| Left Cerebellum VIIb/VIII | -43.5 | -53 | -50 | 1.6 | 53 | 66.25 |
| Right Anterior Cerebellum/Vermis 4-5 | 3.5 | -48 | -10 | 1.5 | 52 | 65 |
| Left Cerebellum Crus2 | -30 | -78.5 | -40 | 1.6 | 50 | 62.5 |

**Table 5:** Peak stereotactic coordinates for BSO gray matter increases compared with AMC.

| Region | x | y | z | -log10(p) | voxels | volume (mm³) |
|---|---|---|---|---|---|---|
| | **Frontal Lobe** | | | | | |
| Right Precentral Gyrus/Superior Frontal Gyrus | 22.5 | -10 | 55 | 2.4 | 859 | 1073.75 |
| Left Superior Frontal Gyrus/BA 6 | -1 | 4 | 60 | 2.4 | 718 | 897.5 |
| Left Supplementary Motor Area/Superior Frontal Gyrus | -3 | -7 | 80 | 1.9 | 293 | 366.25 |
| Left Superior Frontal Gyrus | -23 | -9 | 70 | 1.9 | 211 | 263.75 |
| Left Middle Frontal Gyrus | -42 | 15.5 | 45 | 1.6 | 104 | 130 |
| Left Supplementary Motor Area/Superior Frontal Gyrus | -9 | -2 | 70 | 1.9 | 96 | 120 |
| Right Superior Frontal Gyrus | 18 | 33 | 60 | 1.7 | 82 | 102.5 |
| Right Supplementary Motor Area/Superior Frontal Gyrus | 7 | 2.5 | 60 | 1.9 | 80 | 100 |
| Left Middle Frontal Gyrus | -43 | 7 | 55 | 2.1 | 78 | 97.5 |
| Left Precentral Gyrus | -40 | -11.5 | 50 | 1.9 | 76 | 95 |
| Right Supplementary Motor Area/Superior Frontal Gyrus | 10 | 5.5 | 70 | 1.5 | 72 | 90 |
| Left Middle Frontal Gyrus | -44.5 | 32.5 | 40 | 1.7 | 65 | 81.25 |
| Left Precentral Gyrus | -32.5 | -21.5 | 65 | 1.8 | 61 | 76.25 |
| Left Middle Frontal Gyrus | -28.5 | 45 | 25 | 1.6 | 58 | 72.5 |
| Left Precentral Gyrus | -47 | -6.5 | 55 | 1.6 | 55 | 68.75 |
| Left Middle Frontal Gyrus | -40 | 30 | 45 | 1.9 | 51 | 63.75 |
| | **Parietal Lobe** | | | | | |
| Right Superior Parietal Lobe/Postcentral Gyrus | 48 | -44 | 55 | 2.7 | 5346 | 6682.5 |
| Left Postcentral Gyrus | -27.5 | -43.5 | 75 | 2.2 | 276 | 345 |
| Right Precuneus | 16.5 | -68 | 50 | 2.4 | 102 | 127.5 |
| Left Superior Parietal Lobe/Postcentral Gyrus | -19 | -49.5 | 70 | 2.4 | 83 | 103.75 |
| Left Postcentral Gyrus | -26.5 | -30.5 | 75 | 1.8 | 82 | 102.5 |
| Left Postcentral Gyrus | -49.5 | -25.5 | 60 | 1.9 | 63 | 78.75 |

| | | | | | | |
|---|---|---|---|---|---|---|
| Left Postcentral Gyrus/Precentral Gyrus | -38.5 | -26 | 70 | 1.8 | 63 | 78.75 |
| | **Temporal Lobe** | | | | | |
| Right Middle Temporal Gyrus | 62.5 | -43.5 | -10 | 1.7 | 131 | 163.75 |
| Left Middle Temporal Gyrus | -40.5 | -65 | 20 | 1.5 | 51 | 63.75 |
| | **Occipital Lobe** | | | | | |
| Right Calcarine Fissure | 13.5 | -71.5 | 10 | 2.7 | 178 | 222.5 |
| Right Middle Occipital Gyrus | 50.5 | -70.5 | 0 | 2.1 | 63 | 78.75 |

**Table 6:** Peak stereotactic coordinates for BRCA-preBSO gray matter increases compared with AMC.

| Region | x | y | z | -log10(p) | voxels | volume (mm$^3$) |
|---|---|---|---|---|---|---|
| | **Frontal Lobe** | | | | | |
| Right Middle Frontal Gyrus/BA 10 | 38.5 | 50.5 | -20 | 2.7 | 3198 | 3997.5 |
| Right Superior Frontal Gyrus/BA 10/Medial Frontal Gyrus | 28 | 65 | 20 | 2.4 | 2549 | 3186.25 |
| Left Superior Frontal Gyrus/BA 10/Middle Frontal Gyrus | -13.5 | 65 | 15 | 2.4 | 2506 | 3132.5 |
| Left Middle Frontal Gyrus | -20 | 36 | -10 | 2.2 | 384 | 480 |
| Left Superior Frontal Gyrus | -3.5 | 48.5 | 60 | 2.2 | 346 | 432.5 |
| Left Superior Frontal Gyrus | -17.5 | 61.5 | -5 | 2 | 334 | 417.5 |
| Left Superior Frontal Gyrus | -27 | 57.5 | 20 | 1.9 | 234 | 292.5 |
| Left Superior Frontal Gyrus | -18.5 | 50.5 | 45 | 1.9 | 226 | 282.5 |
| Right Superior Frontal Gyrus | 31 | 55.5 | 35 | 1.9 | 197 | 246.25 |
| Left Superior Frontal Gyrus | -40.5 | 42.5 | 25 | 1.6 | 132 | 165 |
| Left Inferior Frontal Gyrus/BA 47/Middle Frontal Gyrus | -49.5 | 45.5 | -15 | 1.6 | 124 | 155 |
| Left Medial Frontal Gyrus | -5 | 58 | 10 | 1.8 | 116 | 145 |
| Left Supplementary Motor Area | -2.5 | 5.5 | 50 | 1.8 | 113 | 141.25 |
| Right Supplementary Motor Area | 3.5 | 4.5 | 50 | 1.7 | 110 | 137.5 |
| Left Middle Frontal Gyrus | -40 | 30 | 45 | 2.1 | 104 | 130 |
| Left Medial Orbitalfrontal Gyrus/BA 10 | -9.5 | 55.5 | -5 | 1.6 | 103 | 128.75 |
| Left Supplementary Motor Area/Medial Frontal Gyrus | -9.5 | 10 | 50 | 1.9 | 97 | 121.25 |
| Left Middle Frontal Gyrus | -33 | 52.5 | 5 | 1.5 | 96 | 120 |
| Left Medial Frontal Gyrus/Rectal Gyrus | -10.5 | 47.5 | -15 | 1.9 | 88 | 110 |
| Left Anterior Cingulate Gyrus/BA 32/Medial Frontal Gyrus | -11.5 | 26 | 35 | 1.9 | 87 | 108.75 |
| Left Superior Frontal Gyrus/BA 8 | -22.5 | 36 | 50 | 1.7 | 81 | 101.25 |
| Right Supplementary Motor Area/Superior Frontal Gyrus | 12 | 6 | 70 | 1.9 | 71 | 88.75 |
| Left Superior Orbitalfrontal Cortex | -11.5 | 49.5 | -20 | 1.5 | 64 | 80 |
| Left Middle Frontal Gyrus/BA 9 | -43.5 | 32.5 | 35 | 1.7 | 61 | 76.25 |

| | | | | | | |
|---|---|---|---|---|---|---|
| Left Superior Frontal Gyrus | -20 | 55.5 | 40 | 1.7 | 61 | 76.25 |
| Left Inferior Frontal Gyrus | -47.5 | 34.5 | 5 | 1.6 | 54 | 67.5 |
| Left Inferior Frontal Gyrus | -43.5 | 42.5 | 5 | 2 | 54 | 67.5 |
| Right Anterior Cingulate Gyrus/ Supplementary Motor Area/BA 24 | 11 | -4 | 50 | 1.4 | 52 | 65 |
| | **Parietal Lobe** | | | | | |
| Left Precuneus | -11 | -67.5 | 40 | 1.6 | 150 | 187.5 |
| | **Cingulate Gyrus** | | | | | |
| Right Middle Cingulate Gyrus | 6 | 13 | 55 | 1.9 | 209 | 261.25 |
| Right Middle Cingulate Gyrus | 7 | 0.5 | 35 | 2.4 | 208 | 260 |
| | **Cerebellum** | | | | | |
| Right Cerebellum VIII | 43.5 | -61 | -60 | 1.7 | 116 | 145 |

**Table 7:** Peak stereotactic coordinates for exploratory whole-brain ANOVA results for comparing across all four study groups.

| | | | | | | | | Mean Gray Matter Volume | | | |
|---|---|---|---|---|---|---|---|---|---|---|---|
| **Lobe** | **Hemisphere** | **Region** | **x** | **y** | **z** | **# voxels** | **-log10(p)** | **AMC** | **BRCA** | **BSO** | **BSO+ERT** |
| Brainstem | Left | Left Midbrain/Thalamus | 0 | -11.5 | -15 | 406 | 1.74 | 42.15 | 40.18 | 58.97 | 57.16 |
| Cerebellum | Left | Left Anterior Cerebellum | -8.5 | -39 | -20 | 347 | 1.59 | 24.75 | 22.31 | 31.97 | 11.48 |
| Cerebellum | Left | Left Posterior Cerebellum | -35.5 | -63.5 | -20 | 59 | 1.52 | 5.21 | 8.52 | 11.96 | 27.72 |
| Cerebellum | Left | Left Posterior Cerebellum | -2 | -66.5 | -25 | 787 | 1.85 | 11.69 | 31.55 | 26.11 | 19.54 |
| Cerebellum | Left | Left Anterior Cerebellum | -34.5 | -56.5 | -25 | 91 | 1.8 | 89.29 | 99.42 | 97.32 | 91.95 |
| Cerebellum | Left | Left Posterior Cerebellum | -31 | -48.5 | -55 | 98 | 1.36 | 44.99 | 60.54 | 26.99 | 43.19 |
| Cerebellum | Left | Left Posterior Cerebellum | -32 | -75 | -50 | 224 | 1.7 | 58.88 | 61.44 | 53.04 | 55.11 |
| Frontal | Left | Left ACC | -10.5 | 36 | 15 | 440 | 1.52 | 83.16 | 62.25 | 66.90 | 76.07 |
| Frontal | Left | Left Inferior Orbital Frontal | -26 | 22.5 | -15 | 143 | 1.92 | 43.57 | 35.32 | 31.42 | 16.32 |
| Frontal | Left | Left Medial Orbital Frontal | -7.5 | 50 | -5 | 151 | 1.49 | 78.13 | 86.97 | 76.12 | 58.30 |
| Frontal | Left | Left Rectal Gyrus | -7 | 32 | -25 | 114 | 1.7 | 77.19 | 81.65 | 79.80 | 62.05 |
| Frontal | Left | Left Superior Frontal Gyrus | -12.5 | 68.5 | 15 | 86 | 1.92 | 48.37 | 54.79 | 27.65 | 35.87 |
| Frontal | Left | Left Middle Frontal Gyrus | -44 | 8.5 | 55 | 62 | 1.74 | 83.31 | 76.60 | 67.45 | 85.71 |
| Frontal | Left | Left Middle Frontal Gyrus | -44.5 | 33 | 40 | 50 | 1.42 | 91.28 | 91.64 | 80.96 | 87.57 |
| Frontal | Left | Left Inferior Orbital Frontal | -39.5 | 39 | -10 | 92 | 1.8 | 13.19 | 20.41 | 13.07 | 25.54 |
| Frontal | Left | Left Precentral Gyrus | -27 | -12.5 | 75 | 60 | 1.32 | 41.67 | 58.55 | 36.64 | 64.92 |
| Frontal | Left | Left Middle Frontal Gyrus | -30.5 | 41.5 | 25 | 525 | 2.7 | 19.21 | 23.46 | 34.82 | 43.03 |
| Frontal | Left | Left Rectal Gyrus | -2.5 | 54.5 | -30 | 191 | 1.8 | 41.00 | 36.21 | 51.45 | 65.00 |
| Frontal | Left | Left Superior Frontal Gyrus | -20 | 47.5 | 35 | 799 | 1.7 | 16.90 | 46.61 | 35.96 | 21.10 |
| Frontal | Left | Left ACC | -0.5 | 25 | -5 | 88 | 1.47 | 14.22 | 33.42 | 25.66 | 14.96 |
| Frontal | Left | Left Middle Frontal Gyrus | -39 | 44 | 15 | 74 | 1.52 | 82.15 | 90.43 | 90.36 | 82.70 |
| Frontal | Left | Left Caudate/ACC | -14 | 27.5 | -5 | 62 | 1.8 | 23.07 | 42.01 | 40.71 | 16.64 |
| Frontal | Left | Left Superior Medial Frontal Gyrus | -10.5 | 38 | 35 | 64 | 1.47 | 78.47 | 87.48 | 94.54 | 89.30 |
| Frontal | Left | Left Inferior Frontal Gyrus | -47 | 42.5 | -5 | 402 | 1.7 | 40.12 | 28.18 | 46.13 | 51.59 |

| | | | | | | | | | | | |
|---|---|---|---|---|---|---|---|---|---|---|---|
| Frontal | Left | Left Middle Frontal Gyrus | -39.5 | 14.5 | 35 | 83 | 1.8 | 86.15 | 76.68 | 86.24 | 83.19 |
| Frontal | Left | Left ACC | -9 | 33.5 | -5 | 53 | 1.28 | 3.14 | 31.20 | 17.99 | 46.50 |
| Insula | Left | Left Insula | -32 | 2.5 | 5 | 513 | 2.1 | 54.46 | 71.46 | 33.43 | 54.72 |
| Limbic | Left | Left Parahippocampus | -17 | -10 | -40 | 68 | 1.47 | 75.64 | 50.71 | 44.87 | 53.22 |
| Limbic | Left | Left Putamen/Amygdala/Parahippocampus | -21 | 5 | -5 | 496 | 1.8 | 24.17 | 26.10 | 43.81 | 35.75 |
| Limbic | Left | Left Lingual Gyrus/Parahippocampus | -8.5 | -43.5 | 0 | 157 | 1.55 | 4.33 | 14.11 | 23.95 | 5.25 |
| Limbic | Left | Left Parahippocampus/Amygdala/Hippocampus | -24 | -8 | -15 | 84 | 1.4 | 42.01 | 32.54 | 40.58 | 51.90 |
| Occipital | Left | Left Middle Occipital Gyrus | -29 | -81 | 5 | 87 | 1.4 | 94.07 | 80.87 | 85.22 | 81.87 |
| Occipital | Left | Left Middle Occipital Gyrus | -30.5 | -93.5 | -5 | 60 | 1.49 | 62.76 | 59.16 | 36.20 | 58.44 |
| Occipital | Left | Left Cuneus | -16.5 | -84 | 15 | 116 | 1.47 | 49.46 | 78.66 | 59.09 | 72.10 |
| Occipital | Left | Left Middle Occipital Gyrus | -27 | -83.5 | 10 | 71 | 1.38 | 7.55 | 18.25 | 11.51 | 22.56 |
| Occipital | Left | Left Fusiform Gyrus | -29.5 | -67.5 | -5 | 93 | 1.66 | 9.11 | 7.62 | 6.83 | 23.19 |
| Occipital | Left | Left Calcarine/Precuneus | -26.5 | -56.5 | 5 | 863 | 2.4 | 11.63 | 9.47 | 4.84 | 28.24 |
| Occipital | Left | Left Inferior Occipital Gyrus | -33.5 | -80.5 | -10 | 108 | 1.42 | 29.20 | 36.01 | 9.60 | 17.38 |
| Occipital | Left | Left Cuneus/Calcarine | -16 | -75 | 10 | 383 | 2.4 | 39.80 | 53.96 | 32.63 | 40.15 |
| Parietal | Left | Left Superior Parietal Lobe | -26.5 | -47 | 60 | 51 | 1.4 | 36.35 | 35.96 | 48.38 | 62.39 |
| Parietal | Left | Left Postcentral Gyrus | -51.5 | -33.5 | 50 | 56 | 1.66 | 45.77 | 62.53 | 69.10 | 37.76 |
| Parietal | Left | Left Inferior Parietal Lobe | -44 | -35 | 40 | 83 | 1.7 | 3.53 | 17.96 | 16.05 | 25.73 |
| Parietal | Left | Left Inferior Parietal Lobe | -54.5 | -45 | 35 | 174 | 1.66 | 13.09 | 10.48 | 3.11 | 25.13 |
| Parietal | Left | Left Postcentral Gyrus | -56.5 | -27 | 30 | 491 | 1.85 | 74.63 | 91.03 | 49.91 | 68.16 |
| Temporal | Left | Left Middle Temporal Lobe | -40.5 | -68 | 20 | 125 | 1.52 | 30.10 | 14.65 | 11.75 | 6.94 |
| Temporal | Left | Left Superior Temporal Lobe | -36 | -13 | -10 | 67 | 1.55 | 85.39 | 85.32 | 94.01 | 98.00 |
| Temporal | Left | Left Planum Polare/Insula | -39.5 | -5 | -15 | 343 | 2 | 8.32 | 20.02 | 25.61 | 15.26 |
| Temporal | Left | Left Inferior Temporal Gyrus | -49.5 | -39 | -20 | 602 | 2 | 99.67 | 82.67 | 75.86 | 98.48 |
| Temporal | Left | Left Superior Temporal Lobe | -55 | -3 | 0 | 286 | 2.22 | 66.13 | 84.73 | 44.99 | 50.39 |
| Temporal | Left | Left Superior Temporal Lobe | -47.5 | -53 | 15 | 123 | 1.85 | 6.53 | 29.50 | 18.27 | 26.45 |
| NA | NA | NA | 0.5 | 6 | -30 | 126 | 1.62 | 99.35 | 91.57 | 94.81 | 110.20 |
| NA | NA | NA | -12.5 | -14 | -40 | 58 | 1.55 | 61.12 | 70.00 | 78.68 | 40.53 |

| | | | | | | | | | | | |
|---|---|---|---|---|---|---|---|---|---|---|---|
| Brainstem | Right | Right Pons/Medulla | 5.5 | -21 | -50 | 147 | 1.49 | 45.06 | 32.84 | 32.65 | 20.55 |
| Brainstem | Right | Right Pons | 3.5 | -25.5 | -40 | 87 | 1.4 | 44.93 | 37.37 | 51.52 | 18.93 |
| Brainstem | Right | Right Medulla | 11.5 | -35 | -60 | 61 | 1.44 | 10.32 | 22.89 | 6.29 | 9.62 |
| Brainstem | Right | Right Brainstem | 2.5 | -37.5 | -60 | 88 | 1.74 | 4.34 | 19.26 | 20.78 | 3.43 |
| Cerebellum | Right | Right Posterior Cerebellum | 4 | -80.5 | -20 | 566 | 1.49 | 35.96 | 19.36 | 16.12 | 10.94 |
| Cerebellum | Right | Right Posterior Cerebellum | 30.5 | -79.5 | -35 | 551 | 2.4 | 33.92 | 26.64 | 3.93 | 24.56 |
| Cerebellum | Right | Right Posterior Cerebellum | 24.5 | -59.5 | -55 | 1225 | 2.4 | 77.10 | 92.91 | 73.42 | 89.75 |
| Cerebellum | Right | Right Posterior Cerebellum | 39 | -66 | -30 | 52 | 1.38 | 23.97 | 57.23 | 38.83 | 53.19 |
| Cerebellum | Right | Right Posterior Cerebellum | 31 | -85 | -25 | 1106 | 1.92 | 46.78 | 53.61 | 64.97 | 88.61 |
| Cerebellum | Right | Right Posterior Cerebellum | 23.5 | -68.5 | -15 | 330 | 1.47 | 62.83 | 76.09 | 85.42 | 86.24 |
| Cerebellum | Right | Right Posterior Cerebellum | 15.5 | -52 | -35 | 69 | 1.38 | 82.70 | 91.86 | 97.83 | 87.03 |
| Cerebellum | Right | Right Posterior Cerebellum | 39 | -54 | -60 | 481 | 1.92 | 42.23 | 35.44 | 38.98 | 49.70 |
| Cerebellum | Right | Right Posterior Cerebellum | 26.5 | -70.5 | -55 | 51 | 1.34 | 42.20 | 47.22 | 45.14 | 75.39 |
| Cerebellum | Right | Right Posterior Cerebellum | 41.5 | -68.5 | -55 | 51 | 1.24 | 36.23 | 38.57 | 33.84 | 57.69 |
| Cerebellum | Right | Right Posterior Cerebellum | 43 | -50 | -55 | 149 | 1.55 | 5.49 | 8.61 | 21.23 | 11.27 |
| Frontal | Right | Right ACC | 1 | 31.5 | 10 | 857 | 2 | 29.18 | 22.68 | 18.66 | 6.55 |
| Frontal | Right | Right Rectal Gyrus/Medial Frontal Lobe | 6 | 24 | -15 | 106 | 1.52 | 24.16 | 6.91 | 7.26 | 12.96 |
| Frontal | Right | Right Inferior Frontal Gyrus | 33 | 27 | 25 | 70 | 1.44 | 55.34 | 50.65 | 44.66 | 67.79 |
| Frontal | Right | Right Middle Frontal Gyrus | 49 | 36 | 35 | 98 | 1.4 | 35.49 | 62.22 | 50.43 | 31.71 |
| Frontal | Right | Right Superior Frontal Gyrus | 18.5 | 21.5 | 50 | 58 | 1.28 | 49.71 | 61.48 | 90.22 | 44.75 |
| Frontal | Right | Right Middle Frontal Gyrus | 27.5 | 41.5 | 40 | 51 | 1.32 | 75.70 | 59.02 | 59.00 | 76.39 |
| Frontal | Right | Right Middle Frontal Gyrus | 23 | -2.5 | 50 | 2812 | 2.7 | 28.89 | 43.34 | 52.60 | 75.62 |
| Frontal | Right | Right Middle Frontal Gyrus | 32 | 35 | 45 | 161 | 1.8 | 6.86 | 16.12 | 24.54 | 19.04 |
| Frontal | Right | Right Middle Frontal Gyrus | 32 | 7 | 45 | 109 | 1.59 | 6.71 | 12.37 | 11.03 | 38.62 |
| Frontal | Right | Right ACC/Medial Frontal | 13.5 | 43.5 | 20 | 53 | 1.38 | 4.78 | 9.28 | 15.39 | 17.85 |
| Frontal | Right | Right Precentral Gyrus | 40 | -10 | 30 | 198 | 1.74 | 68.57 | 71.76 | 67.72 | 78.42 |
| Frontal | Right | Right Superior Frontal Gyrus | 15.5 | 43 | 35 | 361 | 1.55 | 27.04 | 25.27 | 15.06 | 47.73 |
| Frontal | Right | Right Superior Frontal Gyrus | 22.5 | 36.5 | 60 | 147 | 1.7 | 94.18 | 91.70 | 85.16 | 107.13 |

| | | | | | | | | | | | |
|---|---|---|---|---|---|---|---|---|---|---|---|
| Frontal | Right | Right Middle Frontal Gyrus | 44.5 | 23.5 | 30 | 257 | 1.8 | 38.45 | 40.64 | 17.94 | 21.02 |
| Frontal | Right | Right Superior Frontal Gyrus | 14.5 | 38 | 50 | 227 | 1.59 | 26.94 | 37.22 | 19.09 | 15.34 |
| Frontal | Right | Right Rectal Gyrus | 3 | 54 | -25 | 185 | 1.62 | 7.21 | 25.56 | 18.76 | 20.99 |
| Frontal | Right | Right Middle Frontal Gyrus | 39.5 | 29.5 | 50 | 183 | 1.7 | 61.59 | 60.05 | 41.65 | 38.73 |
| Frontal | Right | Right Inferior Frontal Gyrus | 47 | 16 | 0 | 68 | 1.66 | 91.22 | 66.10 | 93.97 | 99.24 |
| Frontal | Right | ACC | 1 | 34.5 | 5 | 87 | 1.7 | 16.76 | 19.86 | 48.16 | 14.43 |
| Frontal | Right | Right Superior Frontal Gyrus | 10.5 | 23 | 60 | 62 | 1.32 | 10.91 | 4.39 | 23.56 | 11.71 |
| Frontal | Right | Right ACC/Medial Frontal | 1 | 21.5 | -20 | 215 | 1.49 | 36.52 | 58.80 | 25.25 | 37.80 |
| Frontal | Right | Right Precentral Gyrus | 57 | -1.5 | 10 | 55 | 1.34 | 46.14 | 74.57 | 29.72 | 45.95 |
| Frontal | Right | Right Superior Midline | 7 | -53 | 55 | 21065 | 2.7 | 81.41 | 82.21 | 85.93 | 86.22 |
| Frontal | Right | Right Superior/Middle Frontal Gyrus | 44.5 | 56.5 | 10 | 6596 | 2.7 | 1.32 | 25.97 | 17.49 | 21.20 |
| Frontal | Right | Right Superior Frontal Gyrus | 22 | 28 | 55 | 260 | 1.7 | 62.62 | 49.32 | 62.10 | 53.15 |
| Frontal | Right | Right Middle Frontal Gyrus | 34.5 | 21.5 | 40 | 74 | 1.21 | 93.85 | 100.66 | 96.31 | 95.99 |
| Frontal | Right | Right Medial Frontal Gyrus/ACC | 13.5 | 40 | 0 | 58 | 1.4 | 49.29 | 22.48 | 67.98 | 60.07 |
| Insula | Right | Right Insula | 39 | -1 | -15 | 684 | 2.4 | 65.15 | 89.74 | 56.15 | 81.69 |
| Insula | Right | Right Insula | 34.5 | 7 | 10 | 57 | 1.36 | 8.76 | 8.72 | 2.88 | 25.32 |
| Insula | Right | Right Insula | 34.5 | 12.5 | 5 | 263 | 1.92 | 49.56 | 37.18 | 54.32 | 59.82 |
| Insula | Right | Right Insula | 37.5 | -20.5 | 15 | 61 | 1.36 | 55.14 | 70.60 | 53.79 | 53.99 |
| Limbic | Right | Right Parahippocampus | 24 | -21 | -30 | 85 | 1.4 | 58.19 | 39.70 | 35.10 | 45.22 |
| Limbic | Right | Right Parahippocampus/Hippocampus/Amygdala | 35.5 | -13 | -20 | 704 | 1.52 | 77.57 | 42.53 | 73.88 | 42.37 |
| Limbic | Right | Right Hippocampus | 20 | -36.5 | 5 | 62 | 1.3 | 34.03 | 23.89 | 19.55 | 44.05 |
| Limbic | Right | Right Parahippocampus | 21.5 | 7.5 | -20 | 171 | 1.74 | 16.43 | 40.11 | 11.25 | 27.68 |
| Occipital | Right | Right Fusiform/Middle Occipital Gyrus | 30 | -79 | -25 | 85 | 1.36 | 95.61 | 82.70 | 92.36 | 85.07 |
| Occipital | Right | Right Cuneus/Superior Occipital Lobe | 21 | -80 | 15 | 190 | 1.7 | 60.52 | 61.14 | 37.34 | 64.32 |
| Occipital | Right | Right Inferior Occipital Lobe | 46 | -70 | -15 | 251 | 2.4 | 49.57 | 60.06 | 44.99 | 38.74 |
| Occipital | Right | Right Cuneus | 5 | -87 | 10 | 206 | 2.7 | 21.98 | 58.66 | 38.45 | 63.74 |
| Occipital | Right | Right Midde Occipital Lobe | 35 | -70.5 | 35 | 200 | 2.7 | 35.00 | 36.24 | 42.59 | 63.13 |
| Occipital | Right | Right Inferior Midline | 21 | -45.5 | 5 | 7368 | 2.7 | 13.98 | 25.40 | 36.39 | 5.28 |

| | | | | | | | | | | | |
|---|---|---|---|---|---|---|---|---|---|---|---|
| Occipital | Right | Right Inferior Occipital Lobe | 31.5 | -88.5 | -10 | 50 | 1.55 | 52.59 | 34.42 | 34.05 | 55.20 |
| Occipital | Right | Right Cuneus | 16 | -85.5 | 20 | 238 | 1.85 | 42.28 | 59.50 | 66.15 | 63.46 |
| Occipital | Right | Right Calcarine | 31 | -60 | 5 | 55 | 1.52 | 94.11 | 88.21 | 97.08 | 101.06 |
| Occipital | Right | Right Lingual Gyrus | 20 | -64 | -10 | 328 | 1.62 | 4.82 | 24.09 | 19.44 | 31.48 |
| Parietal | Right | Right Superior Parietal Lobe | 24 | -63 | 50 | 372 | 2.1 | 63.25 | 24.37 | 47.02 | 57.45 |
| Parietal | Right | Right Postcentral Gyrus | 39 | -37 | 55 | 74 | 1.36 | 89.94 | 96.89 | 80.70 | 94.52 |
| Parietal | Right | Right Superior Parietal Lobe | 16.5 | -70 | 50 | 59 | 1.4 | 84.26 | 81.13 | 69.04 | 83.51 |
| Parietal | Right | Right Inferior Parietal Lobe | 38.5 | -49.5 | 50 | 2070 | 1.85 | 4.52 | 16.72 | 20.96 | 5.83 |
| Parietal | Right | Right Superior Parietal Lobe | 13.5 | -55 | 70 | 71 | 1.34 | 35.16 | 50.19 | 61.65 | 39.31 |
| Parietal | Right | Right Angular Gyrus/Inferior Parietal Lobe | 46 | -58 | 45 | 149 | 1.38 | 14.83 | 8.21 | 3.60 | 25.11 |
| Parietal | Right | Right Posterior Cingulate/Precuneus | 8 | -53 | 15 | 88 | 1.52 | 52.42 | 51.60 | 44.81 | 61.03 |
| Parietal | Right | Right Angular Gyrus | 48 | -48.5 | 30 | 59 | 1.3 | 58.29 | 33.02 | 64.11 | 75.26 |
| Subcortical | Right | Right Caudate | 15.5 | 28.5 | -5 | 301 | 1.74 | 75.04 | 63.35 | 58.88 | 92.38 |
| Temporal | Right | Right Inferior Temporal Lobe | 50 | -62 | -10 | 58 | 1.44 | 30.98 | 40.50 | 38.79 | 42.54 |
| Temporal | Right | Right Middle Temporal Gyrus | 49.5 | -64.5 | 15 | 455 | 2.4 | 65.49 | 40.51 | 64.95 | 67.54 |
| Temporal | Right | Right Superior Temporal Lobe | 43 | -30.5 | 10 | 53 | 1.3 | 69.85 | 80.63 | 45.97 | 70.59 |
| Temporal | Right | Right Inferior Temporal Lobe | 39.5 | -63.5 | -10 | 102 | 1.7 | 7.52 | 26.27 | 0.91 | 16.22 |
